# Role of Seasonal Importation and Random Genetic Drift on Selection for Drug-Resistant Genotypes of *Plasmodium falciparum* in High Transmission Settings

**DOI:** 10.1101/2023.10.20.563204

**Authors:** Robert J. Zupko, Joseph L. Servadio, Tran Dang Nguyen, Thu Nguyen-Anh Tran, Kien Trung Tran, Anyirékun Fabrice Somé, Maciej F. Boni

**Affiliations:** Center for Infectious Disease Dynamics, Department of Biology, Pennsylvania State University, University Park, PA, USA; Institut de Recherche en Sciences de la Santé, Direction Régionale de l’Ouest, Bobo Dioulasso, Burkina Faso; Nuffield Department of Medicine, University of Oxford, Oxford, UK

## Abstract

Historically *Plasmodium falciparum* has followed a pattern of drug resistance first appearing in low transmission settings before spreading to high transmission settings. Several features of low-transmission regions are hypothesized as explanations: higher chance of symptoms and treatment seeking, better treatment access, less within-host competition among clones, and lower rates of recombination. Here, we test whether importation of drug-resistant parasites is more likely to lead to successful emergence and establishment in low-transmission or high-transmission periods of the same epidemiological setting, using a spatial, individual-based stochastic model of malaria and drug-resistance evolution calibrated for Burkina Faso. Upon controlling for the timing of importation of drug-resistant genotypes and examination of key model variables, we found that drug-resistant genotypes imported during the low transmission season were, (1) more susceptible to stochastic extinction due to the action of random genetic drift, and (2) more likely to lead to establishment of drug resistance when parasites are able to survive early stochastic loss due to drift. This implies that rare importation events are more likely to lead to establishment if they occur during a high-transmission season, but that constant importation (e.g., neighboring countries with high levels of resistance) may produce a greater risk during low-transmission periods.

## Introduction

Despite recent advances in malaria control resulting in a reduction of prevalence, *Plasmodium falciparum* malaria continues to be a major public health concern. The widespread use of artemisinin-based combination therapies (ACTs) has contributed to this reduction in prevalence, but increased usage of ACTs also increases the selective pressure on the parasites to develop drug resistance. Historically, the emergence of drug resistance has followed a pattern of first appearing in low transmission settings, such as Southeast Asia and South America, followed by later migration to high transmission settings. This was the case for chloroquine and sulfadoxine-pyrimethamine resistant *P. falciparum* [1–4], and more recently, artemisinin-resistant *P. falciparum* phenotypes which were identified in western Cambodia in 2007-2008 [5,6]. Since the identification of resistance-associated *kelch13* point mutations [5], artemisinin resistance has been identified in other parts of Southeast Asia [7,8], Guyana [9,10], Rwanda [11,12], and Uganda [13–15]. Thus, developing a mechanistic understanding as to the cause of delayed emergence or slower evolution of drug resistance in high-transmission settings is particularly germane in the African context where a reservoir of *kelch13* mutations currently exists and has the potential for rapid expansion [16].

Several mechanisms have been proposed to explain this pattern of slower drug-resistance emergence, establishment, and/or evolution in higher transmission regions. First, due to higher population-level immunity to malaria, a new infectious mosquito bite is less likely to lead to malaria symptoms in a higher transmission region, resulting in a lower probability that a new infection will be treated by drugs [17–19]. Second, treatment coverage and access are generally lower in high-transmission regions, meaning that new symptomatic infections will also have a lower chance of facing treatment. Third, multiclonal infections (i.e., infections in which the host is infected with several genetically distinct strains of the parasite) result in within-host competition which may suppress drug-resistant clones due to their cost of resistance or immune-mediated competition [20–23]. Within high transmission settings, such as sub-Saharan Africa, multiclonal infections are common [24], thus creating the relevant conditions for drug-resistant and drug-sensitive parasites to be present in the same host. A counteracting factor of this mechanism is that in the presence of the relevant drug therapy, the demise of drug-sensitive parasites in a multiclonal infection may result in competitive release of the drug-resistant parasite and accelerate its spread [21]. Finally, higher rates of recombination in high transmission regions may act against multigenic drug-resistant genotypes by breaking up beneficial combinations of drug-resistance mutations [25] though much work remains to be done on this question.

A growing body of mathematical models suggests that a combination of within-host competition and immune-mediated symptomology are contributors to the cause of the delayed drug resistance emergence in high transmission settings [18,20,21,26–31]. Of particular note is the work of Bushman et al. [21] who used an individual-based model (IBM) combined with ordinary differential equations to model within-host red blood cells, immune response, and parasite dynamics to explore the role of within-host competition. The study found that resistant genotypes initially have a higher risk of extinction in high transmission settings, but resistance can rapidly spread if extinction is avoided. These findings are supported by Whitlock et al. [20] using a similar IBM approach; however, their model also accounted for variations in the antigenic response to various strains of the parasite. Similarly, Masserey et al. [31] also used an IBM coupled with an emulator based approach to examine the impact that various factors such as drug pharmacokinetics/pharmacodynamics, treatment coverage, parasite biology, and environmental factors have on the establishment of drug resistance.

Despite the complexity of the models that have been developed, the effects of spatial-temporal diversity on *P. falciparum* evolution has not been fully explored [32], and it is unknown in which epidemiological scenarios importation of drug-resistant parasites presents the most risk – a question that we explore here. In the context of the high-transmission regions of sub-Saharan Africa, the malaria burden is not uniformly distributed [33], and a country may contain regions of high and low transmission which may influence the evolutionary environment for resistant genotypes. Another limitation of prior studies is that even high transmission regions can have significant seasonal variation in transmission patterns with periods of comparatively low transmission occurring outside of the peak transmission season. Accordingly, there has been increasing interest in the role that seasonality plays in malaria transmission, with recent studies suggesting that persistent asymptomatic infections allow for dry season survival [34]. Finally, while importation is a known mechanism through which drug-resistance has been introduced into various countries, the actual risk of establishment or fixation post-importation is unknown. A contributing factor is the inherent complication in surveillance efforts to monitor importation. While data are limited, in a retrospective study of 54 international travelers arriving in Italy from 2014 to 2015 with confirmed cases of *P. falciparum* malaria, 9 genetic markers for drug resistant genotypes were detected, suggesting that the rate of importation across national borders may be substantial [35], a finding echoed by an earlier study [36].

In this study we explore a straightforward importation mechanism for the introduction of drug resistance in high-transmission regions, through the application of a spatial, IBM of malaria that was previously calibrated and validated for Burkina Faso [37]. This simulation also allows us to explore the role of seasonality and how importation of drug-resistant parasites may lead to the emergence and establishment of drug resistance in a realistic high-transmission context with seasonal variation and heterogeneity of malaria transmission. By restricting importation in the simulation to a particular month, we are able to calculate extinction probabilities and follow long-term trajectories to determine what times of year (and what importation rates) pose the most risk for the establishment and spread of drug resistance.

## Methods

Here we use the term *appearance* to refer to the period following the importation of one or more artemisinin-resistant genotypes (called 580Y for short, using the most common allele found so far). Not all importations are successful, and an imported genotype may immediately go extinct (i.e., no further transmission), or have a brief period of transmission before going extinct. If the genotype is able to survive the action of random genetic drift surrounding its appearance, we say that is has *successfully emerged* once its allele frequency is greater than 0.001 (10^-3^) allowing for possible progression to fixation as the dominant strain [38].

### Simulation Overview

We utilized a previously calibrated and validated spatial IBM of malaria and human movement in Burkina Faso [37,39,40]. As the simulation was designed and constructed with the intent of exploring the evolution of drug resistance in *P. falciparum* [39], it has the appropriate components necessary to explore the mechanisms for delayed emergence without being constructed explicitly for it. The simulation models Burkina Faso as a grid of 10,936 5km-by-5km cells (approximately 273,400 km^2^) and uses an initial population of 3.6 million individuals (25% of the 2007 population) (Supplemental Material 1, §2 – 4). Malaria transmission follows the holoendemic patterns of Burkina Faso with the median *Pf*PR*2-*10 in a given cell ranging between 7.9% to 67.6% [41]. Transmission also follows a seasonal pattern, with transmission increasing at the start of the rainy season in late-May to early-June, peaking between August to October, and declining to a seasonal low in November [37]. On a regional level, the transmission season may be shorter or longer depending upon the length of the rainy season, with the northern Sahelian climatic region having a heighted transmission of about three months, while the season lasts for about five months in the southern Sudanian climatic region.

Upon model initialization, the simulated landscape initially consists of parasites that are chloroquine resistant, artemisinin and piperaquine sensitive, and either amodiaquine sensitive or resistant with a 50-50 probability. Following model burn-in, mutation by the parasite in the presence of the relevant therapy is enabled based upon previously calibrated mutation rates [42], under the assumption that the large majority of mutation occurs during asexual blood stage replication [43]. When locally transmitted infections occur via new infectious bites, individuals may remain asymptomatic, or progress to clinical symptoms based upon their individual immune response. Upon presenting with clinical symptoms, individuals seek treatment following preconfigured rates for Burkina Faso. Individuals under 5 years of age follow treatment-seeking rates determined by previous Malaria Indicator Survey findings [44], while individuals over 5 seek treatment at a rate that is 55% (absolute value) of the under-5 rate. This treatment seeking rate also increases at a rate of 3% starting in model year 2019, consistent with the expected expansion of treatment seeking by individuals [45]. Treatment seeking does not ensure that an efficacious ACT will be taken as the private market accounts for 16.8% of treatments in the simulation, consistent with the local treatment landscape [44].

The individual immune response is summarized as follows, with the full scope elucidated in Nguyen et al. [39] and relevant changes included below. Upon being selected by the simulation to be bitten by an infectious mosquito, the individual undergoes a sporozoite challenge during which their immune response may result in sporozoites being cleared before the parasite enters the liver stage [37]. Based upon the individual’s immune response the probability of infection ranges from 20% (high immune response) to 80% (low immune response). Successful infections proceed to the blood stage where the total parasitemia *D*_*R*_, where *R* notes the specific *P. falciparum* clone, is initially set based upon a random uniform draw from the appropriate parasitemia range (e.g., clinical infections will draw from a range of 10^10^ to 10^12^ parasites per microliter of blood).

Clearance of the parasite by the immune system is then based upon the following calculation:

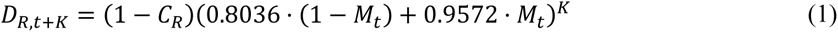

Where *D*_*R*,*t*_ represents the parasite density at time *t*, *M*_*t*_ represents the host immune response using a scale from zero to one, and *CR* represents the fitness cost of the given strain with zero representing the wildtype (i.e., no fitness cost). The parasite density is updated asynchronously every seven days (*K* = 7) and prior validation ensured that this did not differ from daily updating and informed the numeric values used in the equation [39]. In the event of a new infection (resulting in a multiclonal infection), or clearance of a prior infection, the parasite density is updated prior to the next scheduled seven-day update interval.

As individuals are infected and clear infections, the individual immune response; described by the variable *ϴ*; increases according to rates parameterized in Nguyen et al. [39], and impacts the probability that any new infection will progress to clinical symptoms as follows:

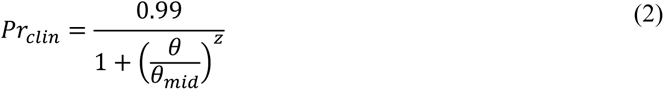

where *ϴmid* describes the point at which immunity confers a 50% chance of developing symptoms, and *z* describes the relationship between the level of immunity and the likelihood of developing symptoms [46]. If individuals go an extended period of time without exposure, *ϴ* decays with a half-life of 400 days, resulting in an increasing likelihood that an individual will experience malaria symptoms following a new infectious mosquito bite.

Individuals are infected in the population based upon the force of infection (FOI) of parasites present in a given cell, with individual host parasitemia used to calculate the FOI of *P. falciparum* clone *R*, at time *t*, for all hosts *n* such that:

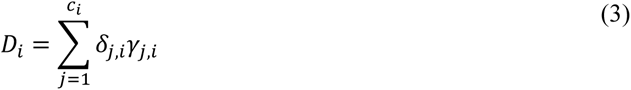

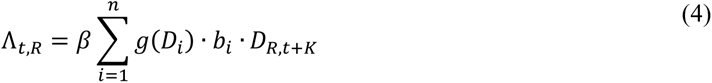

Where Equation 3 describes the total parasitemia *D*_*i*_ of an individual where the quantity of the parasite density of clone *j* in the host is given by *δ*_*j*.*i*_, with *γ*_*j*,*i*_ representing the presence/absence of gametocyte production of the clone *j* where one represents normal production and zero represents none. Due to the nature of the simulation, the residual gametocytaemia following a cured infection (i.e., the asexual parasitemia is zero) is not simulated. To compensate for this the gametocytaemic period is shifted earlier so that infectious hosts have the same number of infectious days as real infections, and overall, this does not impact the simulation since gametocytocidal drugs are not included in the simulation.

Equation 4 describes the FOI for *R* where *bi* is the biting attractiveness of the host, function *g* describes the saturation of transmission probability with increasing parasite density as described by Ross et al. [47], and *β* represents a scaling factor used to calibrate the entomological inoculation rate for the given location within the simulation. Thus, Λ_*t*,*R*_ is ultimately dependent upon the parasitemia of *R* in individuals, resulting in a link to individual immune response and any fitness cost associated with a given parasite. As a result, a high fitness cost incurred by drug resistant strains will result in a discounting of the FOI, consistent with the competitive advantage of the wild type in absence of drug pressure [48,49].

### Study Parameterizations

Using the previously prepared parameterization of Burkina Faso calibrated to the epidemiological situation as of 2017 [37]. While prevalence and case numbers in Burkina Faso are likely to have changed somewhat since 2017, the intent of this study is to explore the general behavior of imported parasites and not exact forecasts of allele frequency. Using this calibration, two simulation studies were conducted, a comprehensive national scale simulation, and a smaller limited locality study. For the first study, the role of seasonality on importation was examined by controlling the month of importation, number of importation events per month (i.e., 1, 3, 6, or 9), and parasite density of the imported infection (i.e., a symptomatic or asymptomatic infection), yielding a total of 96 combinations. During the month of importation, an importation may occur on any day, with the exact number of imports on a given day determined using a Poisson distribution across the entire month. The location of importation is determined by a weighted draw across the entire population, based upon the total population in each cell within the simulation (Supplemental Material 1, §5). Due to the population distribution of Burkina Faso having a lower population in the Sahelian climate zone, this has the effect of biasing importations towards the more densely populated Sudano-Sahelian (containing Ouagadougou, the capital of Burkina Faso) and Sudanian (containing Bobo-Dioulasso, the second largest city in Burkina Faso) climate zones (Supplemental Material 1, §3 – 4.

Upon selection of a cell, the individual to be infected at the location is determined by a uniform random draw from all susceptible individuals at that location. The individual is then infected by an artemisinin resistant *P. falciparum* parasite with the *pfkelch13* allele 580Y, and the host parasitemia level is set to the appropriate value for a symptomatic (between 10^10^ and 10^12^ parasites per microliter of blood) or asymptomatic (less than 1,000 parasites per microliter of blood) infection. In the event of a symptomatic infection, the individual may seek treatment on the basis of their age and the regional treatment seeking rate which ranges from 52.1% to 87.0% for an individual under-5 and 23.4% to 39.1% for an individual over-5. If an individual seeks treatment, they receive either artemether–lumefantrine (68% of treatments), amodiaquine (12.4%), quinine (5.6%), dihydroartemisinin-piperaquine (4.9%), artesunate–amodiaquine (2.5%), artesunate (2.2%), artesunate, sulfadoxine/pyrimethamine (1.6%), chloroquine (1.5%), or mefloquine (1.3%) with the probability of a specific treatment based upon previous survey data and the make-up of the nationally recommended first-line therapies and private-market treatments [37,44,50]. Although importation is implemented by importing a singly-infected individual, following importation multiclonal infections are possible if the individual is infected by another clone. Following model burn-in, the simulation is allowed to run for twenty years. In order to ensure the statistical validity of the results, 50 replicates of each combination were run, for a total of 3,600 replicates.

Following evaluation of a drug-resistant genotype establishing under various importation conditions, additional reporting was incorporated to capture additional population immunity data and three additional parameterizations were prepared. These changes allowed for an assessment of the mechanisms of delayed drug resistance or establishment to be evaluated, within the constraints of the mechanisms that are implemented within the simulation. For the first parameterization, *de novo* mutations were enabled using a previously determined mutation rate [42] so that the evolution and spread of drug-resistant genotypes could be observed

Finally, additional model validation was conducted to ensure conformity of multiclonal infections and multiplicity of infections (MOI) to field conditions. Within the simulation, the proportion of multiclonal infections fluctuates on a seasonal basis, and the ranges are in good agreement with previous studies. A sentinel site study in Nanoro, Burkina Faso, located in the Sudano-Sahelian climate zone, was conducted from September 2010 to October 2012 and recorded a mean MOI of 2.732 (±0.056) with a range of 1 to 7 parasite genotypes [51], compared to simulation results of 2.220 ± 0.455 (Supplemental Material 1, §1).

### Limited Locality Studies

Following completion of the national scale studies, additional studies were conducted with a limited population or limited geographic scope intended to isolate or reproduce dynamics observed in the national-level model results. Specifically, these more limited models were used to explore if observed seasonal fluctuations in 580Y frequency and treatment seeking behavior could help to explain observed transmission and infection dynamics. A total of three additional spatial models were prepared, all deriving from the same configuration used for the national scale model: a single cell with a population of 100,000, a two-by-two grid with a total population of 300,000 individuals, and a three-by-three grid with a total population of 320,000 individuals. All models are based upon four different configurations in which the five-month seasonal pattern of the Sudanian zone is enabled or disabled, and treatment seeking is either balanced (i.e., 50% of under-5, 50% of over-5) or skewed based upon the national upper and lower bounds (i.e., 87% of under-5, 23.4% of over-5).

### Statistical Analysis

To examine whether the month of case importation is associated with the frequency of parasites with the 580Y allele, we performed a Kruskal-Wallis test to identify whether differences exist across months, followed by pairwise Wilcoxon rank-sum tests to identify which pairs of months yielded significantly different frequencies following importation. This was repeated, collapsing months into the high (June – October) and low (November – May) transmission seasons to compare these two time periods. This was followed by examining whether different months of importation are associated with a greater probability of emergence of the 580Y allele, defined by having a frequency ≥ 0.001, we used chi-squared tests for proportions. An initial, global chi-squared test identified if any months differed, and subsequent pairwise tests identified months with different probabilities. Within months, we also tested whether symptomatic or asymptomatic importation was associated with probability of establishment. All p-values lower than 10^-4^ are reported as 10^-4^.

To show alignment among simulated time trends in total infections, treatment administration, immune response, and frequency of parasites with the 580Y allele, Spearman’s correlation coefficients were calculated following visual inspection. For pairs of variables with oscillating trends that did not have aligned peaks and troughs, a lag was applied to align the oscillations, providing insight into how one trend follows another. The use of a lag is appropriate given the inherent delays associated with infection, presentation of symptoms, and transmission of *P. falciparum* infections. Correlation coefficients were calculated for the three climatic regions for scenarios consisting of *de novo* mutation, importation during the low transmission season, and importation during the high transmission season.

## Results

### Role of Seasonality on Importation

When varying the month of importation of drug resistance, the number of importations, and the parasitemia of the imported individual, there is a clear difference between extinction outcomes and sustained transmission outcomes when comparing low-transmission months to high-transmission months (Figure 1). In six of the eight combinations for symptomatic/asymptomatic importation and importation count, drug-resistant genotypes are more likely to establish when imported during low-transmission months (p ≤ 0.0007, Wilcoxon-Rank Sum, Supplemental Materials 2, Table S2). In the scenarios of three asymptomatic importations or one symptomatic importation, future resistance frequencies appear lower for parasites imported during low-transmission periods (Figure 1), but the differences are not statistically significant (*p*=0.81 and *p=*0.41, respectively) due to the large number of zeros in each set of simulations. This large number of zeros for configurations with one or three asymptomatic importations, or one symptomatic importation per month, underlines the influence of extinction and random genetic drift during the importation process. If an imported parasite is unlikely to be sampled by a mosquito, then low transmission periods would be associated with lower risk of establishment. This is most easily seen in the extinction paths in Figure 2 where importation is rare (i.e., one importation event in a given month) and onward transmission occurs with low probability due to the asymptomatic nature of the imported infections; in these scenarios, extinction probabilities are higher for parasites imported during the low-transmission season. However, averaging across all scenarios, pairwise comparisons of months across the low season and the high season indicate that 580Y allele frequency after ten years is likely to be a median 1.84-fold higher (IQR: 0.90 – 3.35) if the allele is imported during the low-transmission season (Supplemental Materials 2, Table S4), indicating that if importation events are common (i.e., several per month) low-transmission periods are associated with a higher starting frequency and a higher a probability of emergence or establishment for the recently imported parasite.

**Figure 1.**
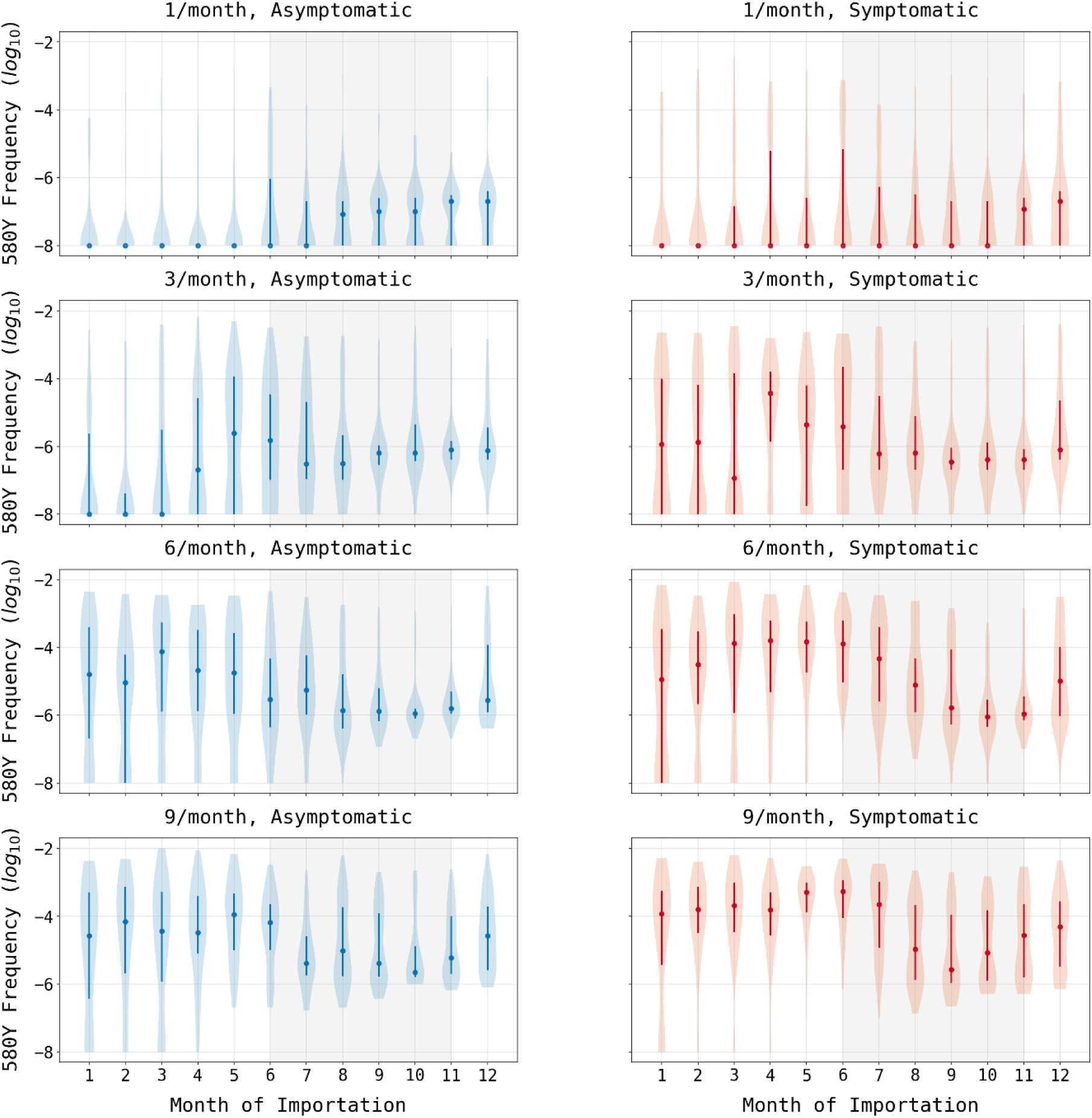
580Y frequency at model completion (after 20 years) based upon month of importation. Circles show median allele frequency, bars show interquartile ranges, and violin plots show full range. As expected, the final frequency of 580Y increases as the number of importations increases (top to bottom) and when cases are symptomatic as opposed to asymptomatic (left to right). In most scenarios, importations that occur during periods of low seasonal transmission are more likely to result in establishment than cases imported during periods of high seasonal transmission (shaded region).

**Figure 2.**
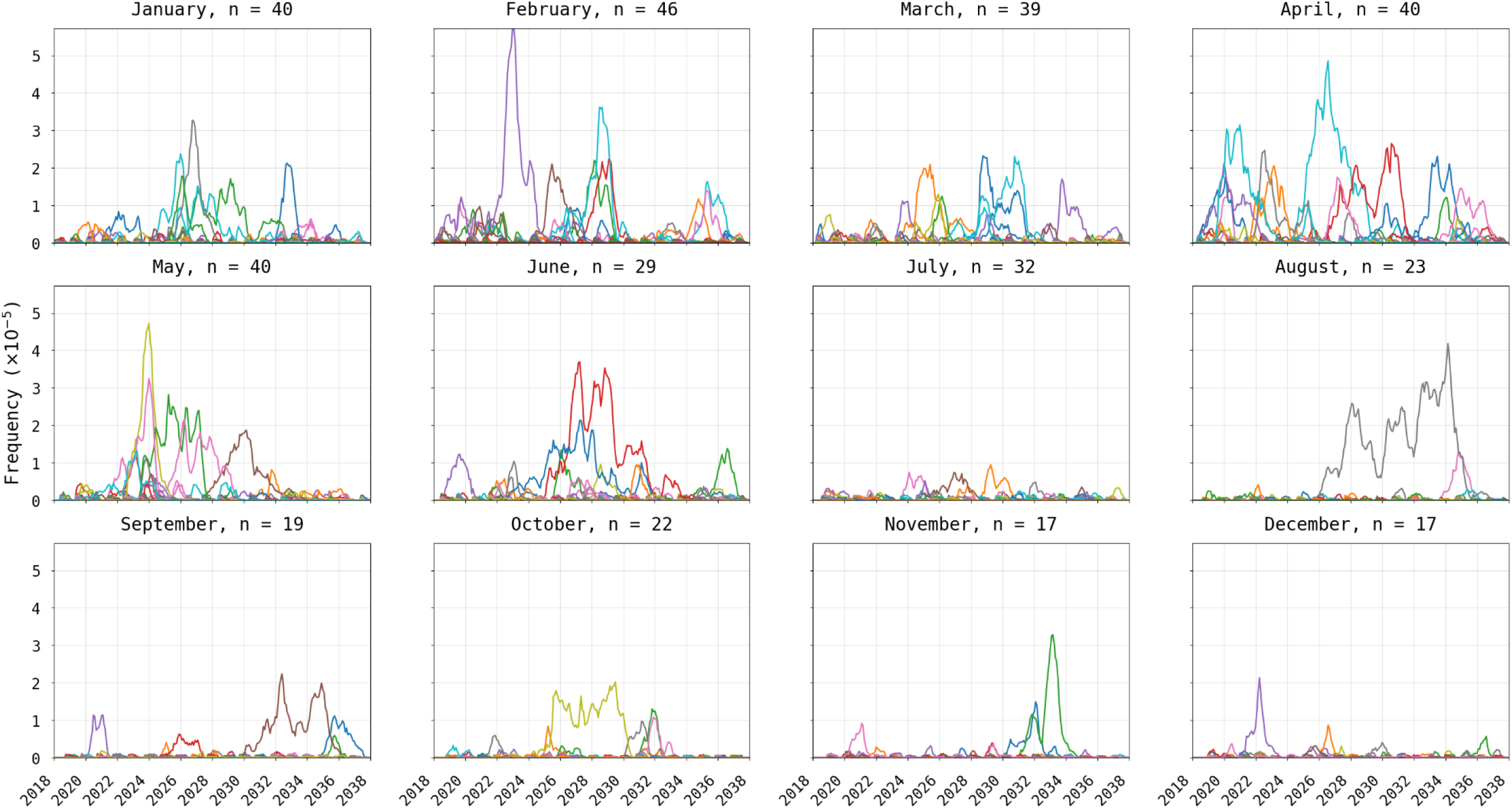
Visualization of 580Y trajectories that reached extinction. Plots show 580Y allele frequency trajectories under a scenario of one asymptomatic importation per month and are broken up into twelve panels by month of introduction. Only trajectories that reached extinction, out of fifty model runs, are shown. Title on each panel shows the month of introduction and the number (*n*) of trajectories that reached extinction in the first 20 years. The likelihood of extinction was higher in the low transmission season (78% to 92% during Jan-May) than in the high-transmission season (34% to 64% during Jun-Dec).

While any imported parasite has a small chance of surviving past initial appearance, the likelihood of progressing to a frequency of 10^-3^, suggesting emergence has occurred and the parasite is likely to be observed, in any scenario was generally below 30% (Figure 3). As expected, our analysis shows that a higher number of importation events is associated with a higher likelihood of eventual establishment (asymptomatic global χ^2^ = 109.3, df = 3, p < 0.01; symptomatic global χ^2^ = 145.9, df = 3, p < 0.01). Median probability of progressing past an allele frequency of 10^-3^ is 0.08 (IQR: 0.02 - 0.14; across 56 month- scenario combinations) when importation occurs during the low-transmission season, and 0.02 (IQR: 0.02 – 0.06; across 40 month-scenario combination) during the high transmission season. These results support the finding that when a drug-resistant genotype is imported, assuming it can escape the risk of random extinction, emergence is more probable for imports during the low-transmission season. When importation is common (9 asymptomatic imports per month, Figure 4), only 2% to 8% of high season importations (i.e., between June and October) reach a 580Y allele frequency ≥10^-3^, whereas 4% to 18% of the low season importations reach a 580Y frequency ≥10^-3^. When the months immediately following the high transmission season are excluded (i.e., November and December), the range is 12% to 18%, suggesting selection mechanisms at play during the high transmission season may are still at play for a period of time following the end of the season.

**Figure 3.**
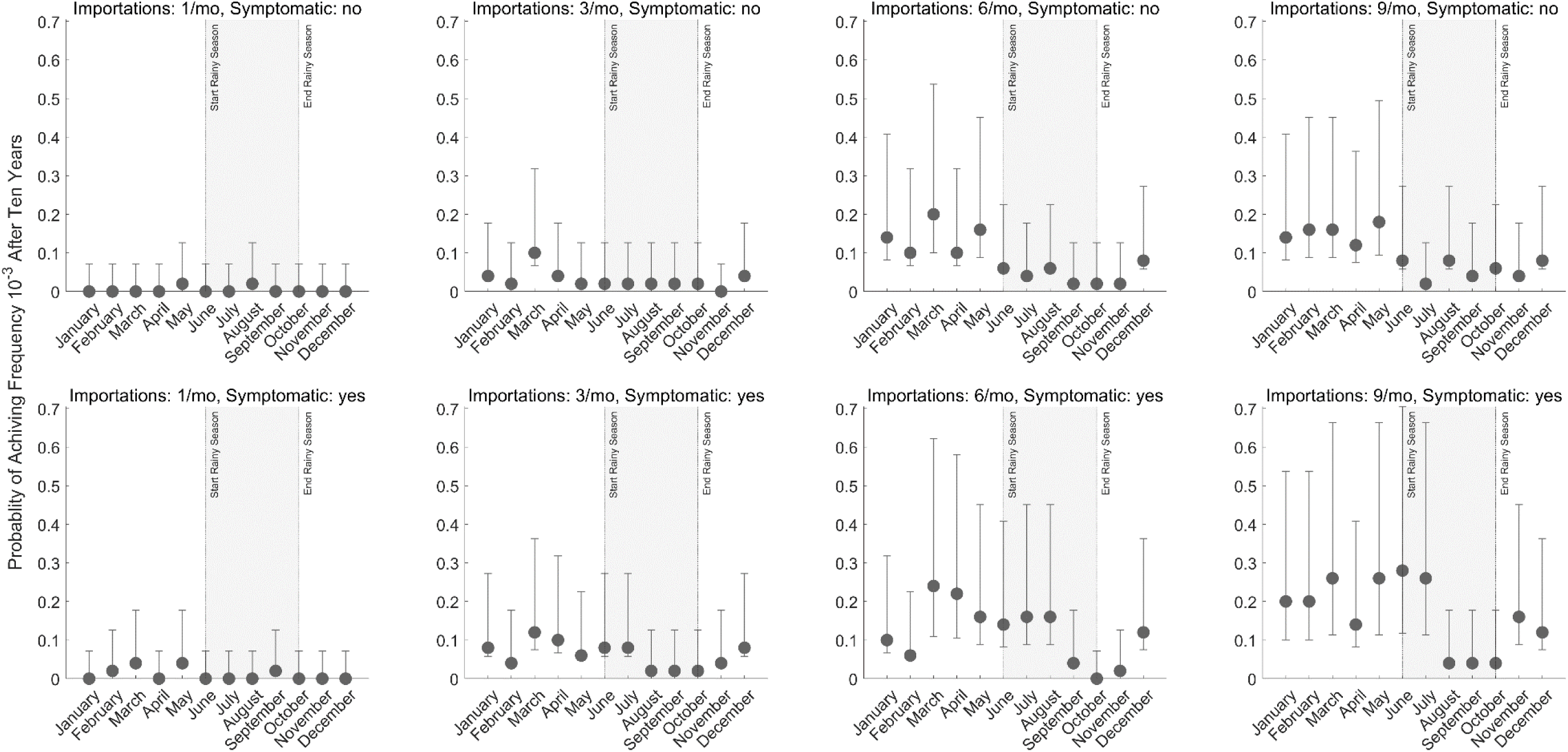
Probability of successful emergence following importation. Probabilities shown (circles) are maximum likelihood estimates from fifty simulations and bars show 95% confidence intervals (exact binomial method). Probabilities of emergence are stratified by month of importation (*x*- axis), by number of importation events per month (columns), and by whether the imported parasite occurred in an asymptomatic (top row) or symptomatic (bottom row) individual. Successful emergence is generally more likely for parasites imported during low transmission season (non- shaded).

**Figure 4.**
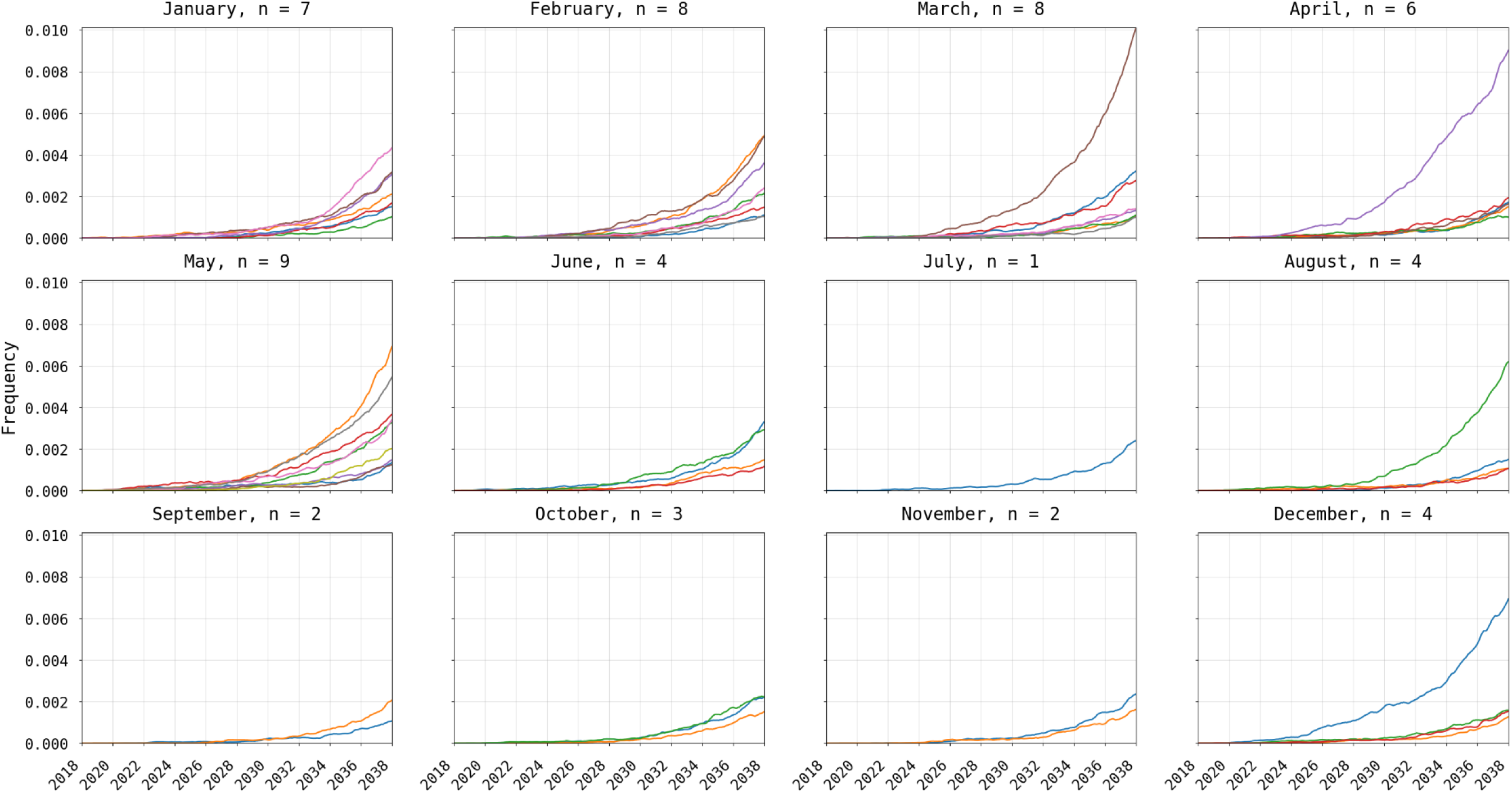
Visualization of 580Y trajectories that successfully emerged (frequency > 0.001). Plots show 580Y allele frequency trajectories under a scenario of nine asymptomatic importations per month and are broken up into twelve panels by month of introduction. Only trajectories that reached an allele frequency >0.001, out of 50 simulations, are shown. Title on each panel shows the month of introduction and the number (*n*) of trajectories that successfully emerged. Successful emergence was higher in the low transmission season (12% to 18% during Jan-May) than in the high-transmission season (2% to 8% during Jun-Dec).

### Selection mechanisms during high- and low-transmission seasons

An examination of the short-term changes in malaria dynamics as the transmission season changes suggests three common effects that may influence the selections strength for drug-resistant genotypes: changes in drug coverage, changes in symptoms occurrence, and changes in the multi-clonal nature of some infections. In a seasonal malaria setting, the age distribution of cases can change between low and high season, and if treatment coverage depends on age, then selection pressure for drug resistance will vary by season as the total population with clinical infections who seek treatment (i.e., total population treatment coverage) varies by season. A simple demonstration of this can be seen in a small population model (320,000 individuals occupying a 3x3 grid) with age-based treatment coverage and seasonal transmission presented as factors in the analysis. When both seasonality (based upon the five-month Sudanian zone in Burkina Faso) and age-based treatment coverage (87% coverage for under-5 and 23.4% coverage for over-5) are present then treatment coverage varies seasonally, in this example between 54% and 60% (Figure 5), in contrast to constant treatment coverage when seasonality is absent or under equal treatment seeking. The under-5 and over-5 treatment coverages in this example were chosen as representational of the maxima and minima for each group based upon provincial coverages. In our national scale simulations of Burkina Faso, population treatment coverage changes by 2% to 3% (absolute value) between the seasons resulting in weak to moderate changes in selection pressure (Figure 6).

**Figure 5.**
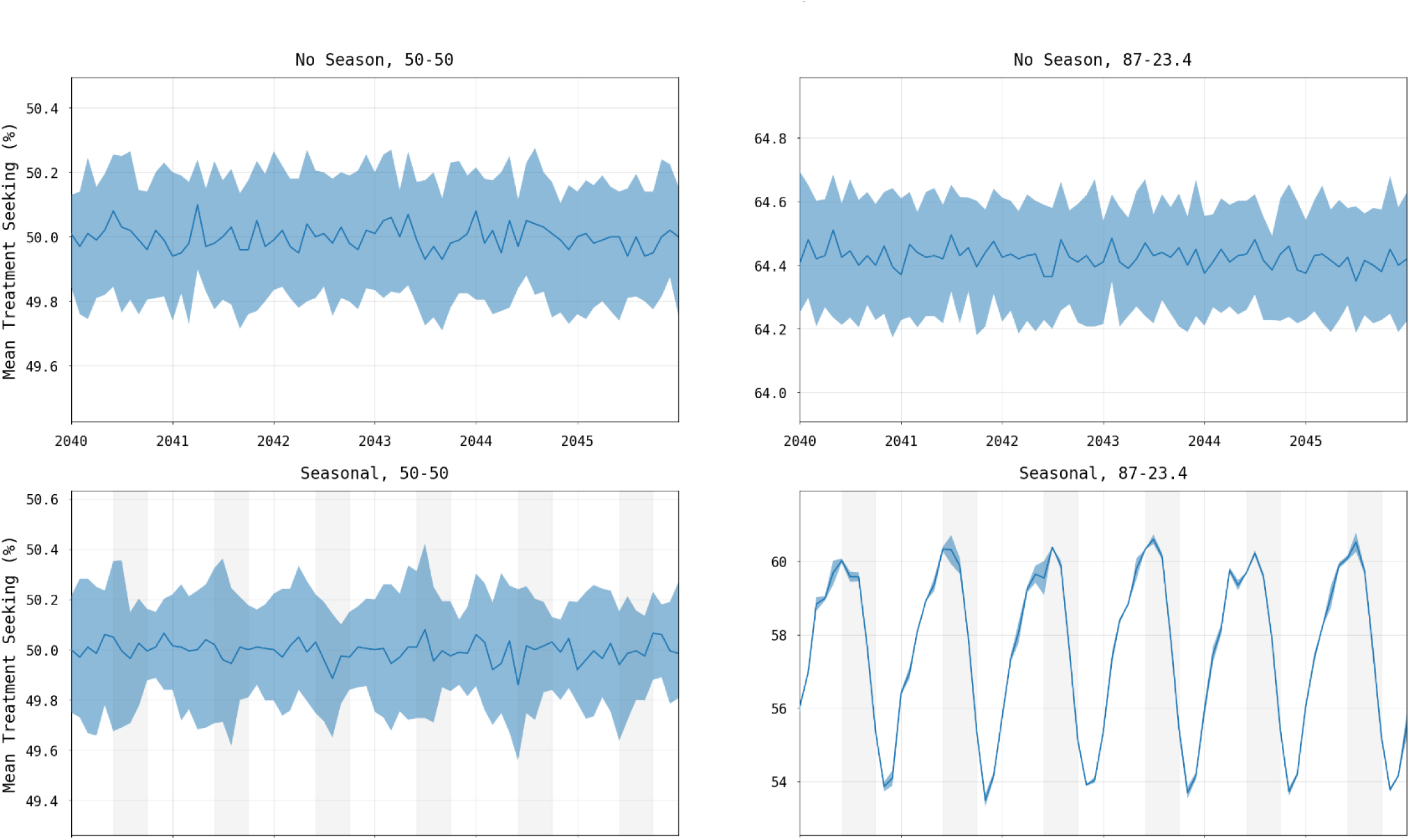
Change in treatment seeking when controlling for seasonality and treatment seeking by age group. Lines in each panel show median percentage of symptomatic malaria infections seeking treatment with shaded areas showing interquartile ranges from one hundred simulations. Panel titles show whether the epidemiological setting represents seasonal (right) or non-seasonal transmission (left), and whether treatment seeking is the same across age groups (“50-50”) with 50% of individuals seeking treatment (left) or uneven with 87% of children under- 5 and 23.4% of individuals over-5 seeking treatment (right). In the presence of both seasonality and uneven treatment seeking across age groups, treatment coverage and thus selection pressure change through time (bottom right).

**Figure 6.**
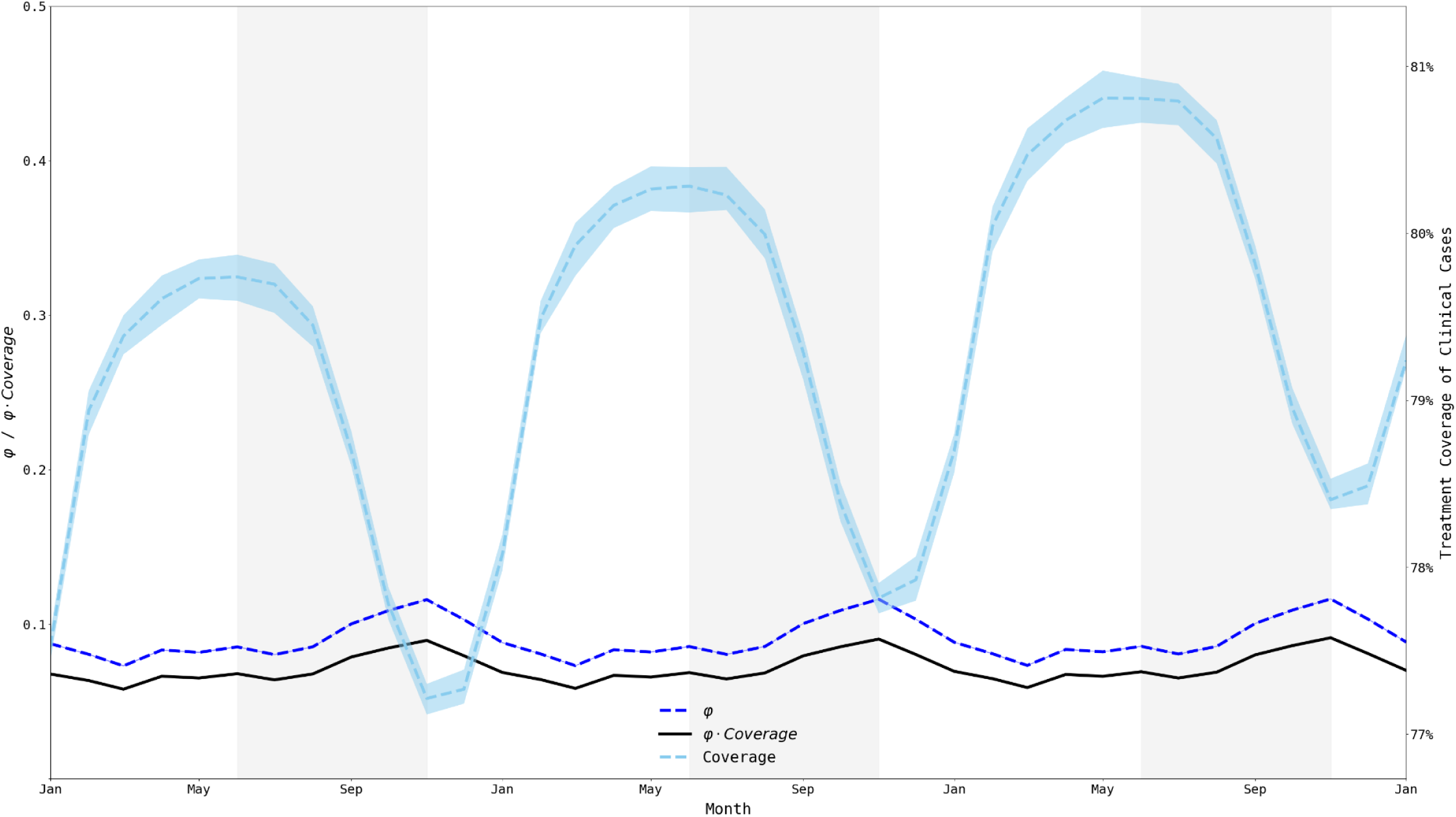
Treatment coverage and fraction symptomatic (*φ*) from Jan 1, 2033, to Jan 1, 2036. When examining a 36-month window, we clearly see that the population treatment coverage is slowly increasing (consistent with a gradual increase in treatment seeking over time) and that treatment coverage (right-axis) fluctuates moderately with the transmission season. The fraction of all infections that are symptomatic (*φ*) remains relatively constant (between 0.073 and 0.117) but fluctuates out-of-phase with treatment coverage. The product of *φ* and coverage (black line) fluctuates between 0.058 and 0.092. Medians and IQRs (shaded areas) shown from fifty simulations.

The nature of the simulation allows for the mean level of all individual immune responses to the parasite to be captured, denoted here as *θ*_*pop*_. As expected, *θ*_*pop*_ follows a lagged seasonal cycle, with *θ*_*pop*_ having the strongest Spearman correlation with number of infections three months prior (Figure 6), which is independent of the number of importations across the six combinations of region and importation timing. This cycle of *θ*_*pop*_ increasing and decreasing across seasons creates a linkage between individual immune responses and the likelihood of symptoms and treatment seeking. However, the *θ*_*pop*_ value peaks and troughs between 0.42 and 0.45 (on an immunity scale of zero to one) warranting further investigation as to whether immunity differences of this magnitude can have an observable effect on selection pressure.

One consequence of *θ*_*pop*_ changing between seasons is that the fraction of malaria parasites currently residing in symptomatic patients versus asymptomatic patients changes as well. The quantity *φ*, defined as the ratio of symptomatic infections to all infections [17,19,52] gives a general description of what proportion of drug-resistant genotypes are currently experiencing positive selection resulting from treatment and what proportion are currently undergoing negative selection imposed by their fitness cost. This symptomatic fraction *φ* appears to be generally low, ranging from 0.07 to 0.12, with a higher proportion of infections subject to drug pressure at the end of the high season than during the middle of the low season. However, *φ* is out of phase with the treatment coverage suggesting that the net combined effect of treatment coverage and symptoms presentation may result in negligible changes in evolutionary pressure during the course of the year (Figure 6).

The role of the individual immune response also works in tandem with the role of within-host competition occurring in multiclonal infections. The high MOI – with a median ranging from 1.747 – 2.268 depending upon the scenario and climatic zone (Supplemental Materials 1, §3) – along with the low frequency of resistant clones during the appearance and emergence of drug-resistant genotypes indicates that within-host competition between drug-sensitive and drug-resistant genotypes is likely occurring. The strength of this effect can be examined within the simulation by comparing the proportion of multiclonal infections carrying a 580Y clone to all multiclonal infections (Figure 7). This proportion is highest when median MOI is lowest, towards the end of the high-transmission season, again showing that these seasonal forces are acting in opposition preventing the formation of a clear picture of when within-host competition against drug-resistant genotypes should be the strongest.

**Figure 7.**
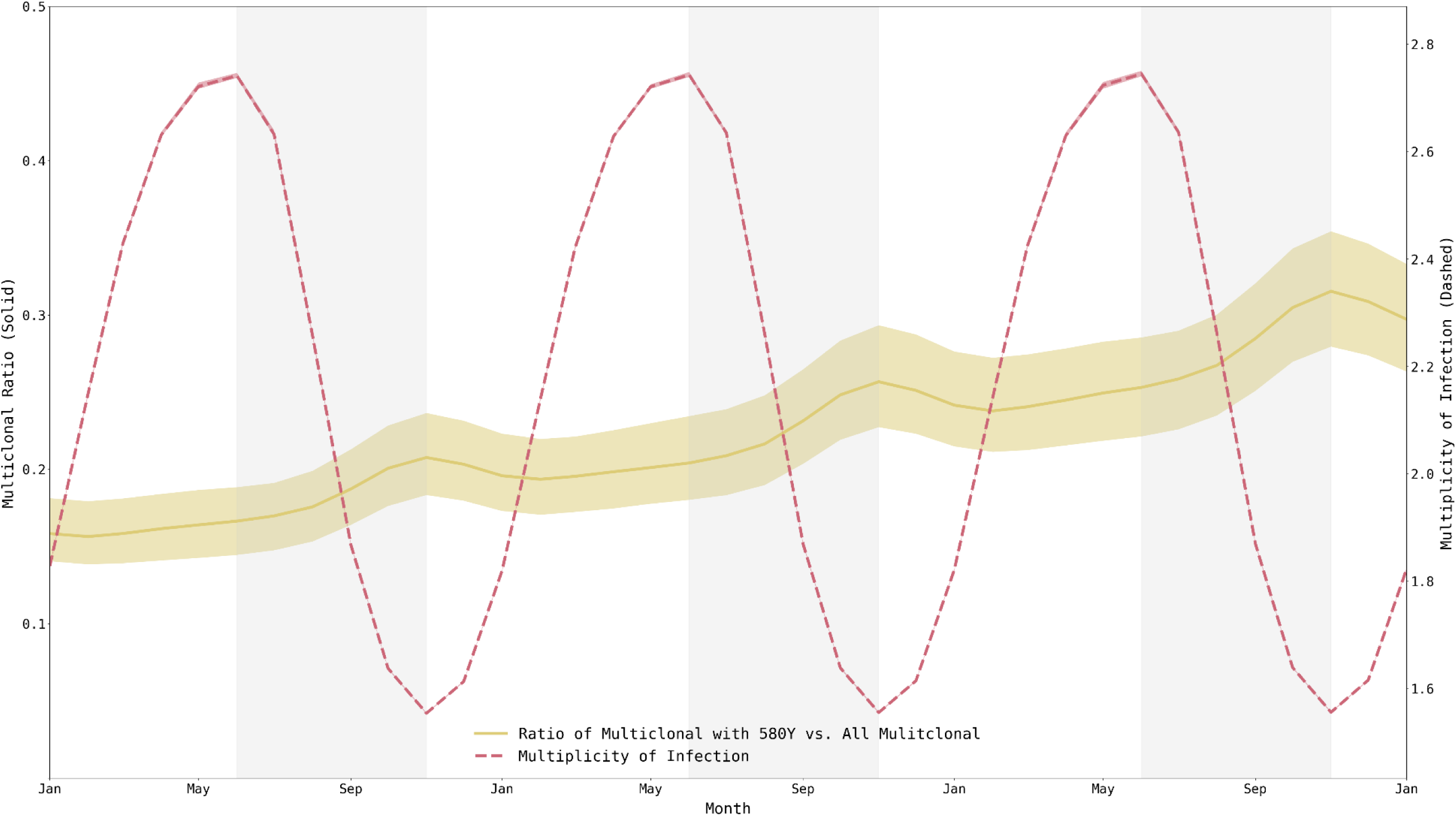
Multiplicity of infection and fraction of multi-clonal infections harboring resistant alleles, from Jan 1, 2033, to Jan 1, 2036. Mean multiplicity of infection (MOI) across individuals ranges from 1.55 to 2.75, peaking at the end of the low-transmission season. Fraction of multiclonal infections that harbor resistant alleles also fluctuates and peaks at the end of the high-transmission season (out of phase with MOI). There does not appear to be a particular period when 580Y alleles are experiencing maximum within-host competition from wild-type parasites. Medians and IQRs (shaded areas) shown from fifty simulations.

## Discussion

Drug resistance has always presented a danger to public health goals. For malaria, however, there is time to prepare as antimalarial drug resistance typically emerges slowly and may take a decade or more to spread geographically. Here, we look at the effects that importation of drug-resistant genotypes has on the national-scale malaria epidemiology (modeled here as the high-transmission settings of Burkina Faso) and we ask whether imported drug-resistant genotypes are more likely to establish if importation occurs in the high-transmission season or in the low-transmission season. We show that random genetic drift and starting frequency play an important role in an imported allele’s future trajectory, but we are uncertain if change in selection pressure is substantial enough across transmission seasons to alter a drug-resistant genotype’s evolutionary path after importation.

The major evolutionary-epidemiological gap identified in our work is that seasonally changing selection pressures for drug resistance are not easily identified as such, and that the direction of change may not always be clear. Three common factors are known to affect drug-resistance evolution across transmission settings – treatment coverage, symptoms presentation, and within-host competition in multi- clonal infections – but these factors do not align the same way between seasons in the same epidemiological setting as they do between countries that have different epidemiological settings. For example, in the seasonal setting presented here, treatment coverage goes up seasonally at the same time as symptoms presentation goes down. The proportion of multi-clonal infections harboring at least one drug-resistant genotype is highest when median MOI is lowest, leading to an ambiguous picture or perhaps small evolutionary differences in terms of when within-host competition may be acting to reduce the frequency or relative density of drug-resistant genotypes.

The traditional population-genetic effects of importation and drift do have their expected behaviors in our analysis of drug-resistance importation for malaria. Random genetic drift may lead to extinction for imported mutants with higher probabilities of extinction associated with low importation rates and low transmission rates. Imported parasites that are not lost due to drift will progress to emergence and establishment more quickly if the initial importation event occurred in a smaller population, i.e., during low-transmission season.

As in all epidemiological modeling analyses, the model structure itself means that some limitations are present in the analysis and interpretation. First, the way that symptomatic importations are implemented may introduce some bias in favor of the parasite. Specifically, (*i*) immune response is ignored when an infection is imported and, (*ii*) treatment seeking behavior is based upon where the individual resides; however, in practice individuals may be more (or less) likely to seek treatment if symptomatic when passing through a port of entry. Second, genetic background and age were also not included as factors in the analysis. Third, this study focused on importation of *pfkelch13* mutants associated with longer clearance half-lives and high rates of treatment failure, but the treatment failure rate of *pfkelch13* mutants depends strongly on the presence/absence of certain partner-drug mutations [53,54], as well as the genetic background that these mutations appeared on. Imported parasites with intrinsically high failure rates would likely have an easier time avoiding the effects of drift, spreading, and establishing in the population. Finally, age, as is well known in malaria, is an important factor associated with malaria history and treatment seeking; and an imported parasites in a younger patients will have different parasitemia levels and different likelihood of seeking treatment depending on the patient’s country of origin, recent history in high transmission settings, and malaria immunity.

The critical public health conversation that this study has implications for is the future of molecular surveillance for specific *P. falciparum* drug-resistant genotypes that are at risk of being imported from one country to another. In particular, when transmission is highly seasonal, monitoring for known markers of drug resistance may be more important during the low-transmission season. This may become increasingly relevant in the African context with the *de novo* appearance of the drug resistance markers 561H in Rwanda [11], along with 469Y and 675V in Uganda [14]. In addition to the *de novo* appearance of these markers on a regional level, at least one instance of the 561H allele has been isolated in Uganda [14,15], suggesting that cross border migration of drug resistance is already taking place.

A major factor for projecting the future evolution of drug resistance is the increased usage of ACTs within high transmission settings. While the historical pattern has been for the establishment of imported genotypes following the evolution of drug resistance in low transmission settings, the recent identification of *de novo* drug resistance in high transmission settings [11,13,16] suggests that low-transmission appearance may simply be more likely but not a determinative rule for all drug-resistance emergence events. Given the role that individual immune response plays in creating an environment conducive to the evolution of drug resistance, understanding the possible impact of upcoming vaccinations (i.e., RTS,S/AS01) on selection for – or against – drug resistance by the parasites may play a role in speeding up or slowing down the selection of drug-resistant genotypes [55,56]. Nevertheless, despite some of the known effects of transmission setting on drug-resistance evolution – via differences in drug coverage and symptoms presentation, primarily – these effects do not appear to translate to differences between seasons in the same epidemiological setting. While drift and importation rate do appear to have their traditional effects on the success of recently imported genotypes into a new population, natural selection on drug resistance does not appear to be stronger in one part of the malaria season than another, or these selective differences could not be identified in the Burkina Faso-specific model parameterizations analyzed here.

## Availability of data and materials

The source code for the base mathematical model and analysis specific to this manuscript can be found on GitHub at https://github.com/bonilab/malariaibm-spatial-BurkinaFaso-2022. Within the repository the dataset(s) supporting the conclusions of this article (and its additional files) are stored under, https://github.com/bonilab/malariaibm-spatial-BurkinaFaso-2022/tree/main/Data

## Acknowledgements

Simulations described in this study were performed on the Pennsylvania State University’s Institute for Computational and Data Sciences’ Roar supercomputer.

