## Supplemental Material 1 for "Role of Seasonal Importation and Random Genetic Drift on Selection for Drug-Resistant Genotypes of *Plasmodium falciparum* in High Transmission Settings"

Supplemental Materials 1 for “Long-term effects of increased adoption of artemisinin combination therapies in Burkina Faso” Zupko et al. (2023)

### 1 Multiplicity of infection

Table S1. Multiplicity of infection under scenarios used to assess the delayed drug resistance selection.

| **Scenario** | **Climate Zone** | **Mean** | **Median** | **IQR** |
| --- | --- | --- | --- | --- |
| *De Novo* | Sahelian (3 month) | 1.725 ± 0.276 | 1.747 | 1.475 – 2.003 |
|  | Sudano-Sahelian (4 month) | 2.220 ± 0.455 | 2.263 | 1.854 – 2.699 |
|  | Sudanian (5 month) | 2.187 ± 0.499 | 2.028 | 1.671 – 2.649 |
| *Low Transmission* | Sahelian (3 month) | 1.726 ± 0.277 | 1.750 | 1.476 – 2.005 |
|  | Sudano-Sahelian (4 month) | 2.222 ± 0.456 | 2.267 | 1.854 – 2.700 |
|  | Sudanian (5 month) | 2.189 ± 0.500 | 2.032 | 1.673 – 2.657 |
| *High Transmission* | Sahelian (3 month) | 1.726 ± 0.276 | 1.751 | 1.476 – 2.005 |
|  | Sudano-Sahelian (4 month) | 2.222 ± 0.456 | 2.268 | 1.854 – 2.701 |
|  | Sudanian (5 month) | 2.189 ± 0.500 | 2.032 | 1.673 – 2.658 |

The multiplicity of infection (MOI) was calculated using a Python script at the same time as the generation of the plots used in the manuscript. The results shown in Table S1 are based upon all timepoints and replicates following model burn-in. As expected, there is little variance in the MOI between scenarios, but some between the climatic zones as a result of the duration of the high transmission season coupled with the resident population within the zones.

### 2 Figure S1. Population density of Burkina Faso.

Boundary data from World Bank Group (2018), population data from WorldPop (2018).

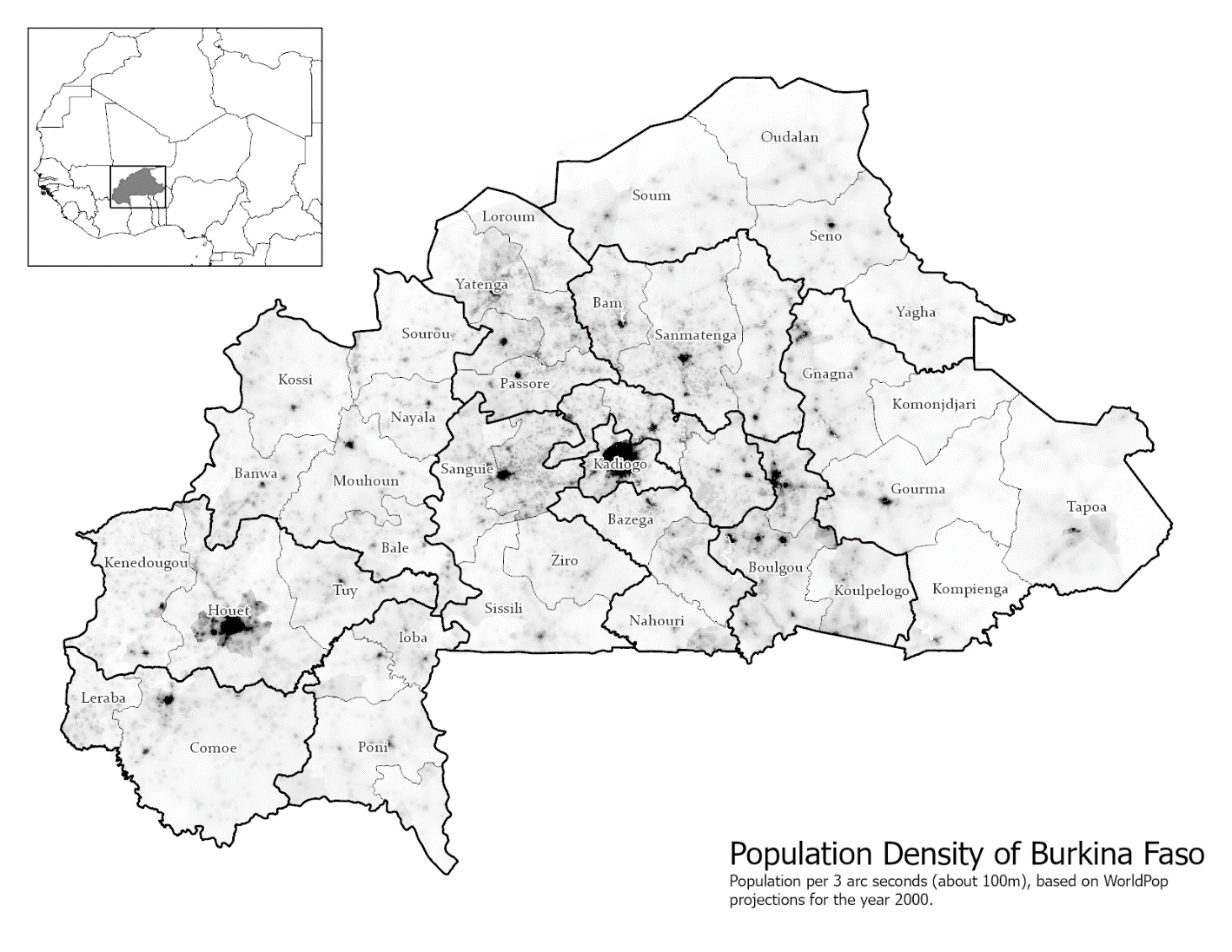

### 3 Figure S2. *Pf*PR_2-10_ in Burkina Faso for 2017

Annual mean projected Plasmodium falciparum prevalence for ages 2 to 10 (*Pf*PR_2-10_) in 2017 (Weiss et al., 2019), boundary data from OpenStreetMap (2017) and World Bank Group (2018).

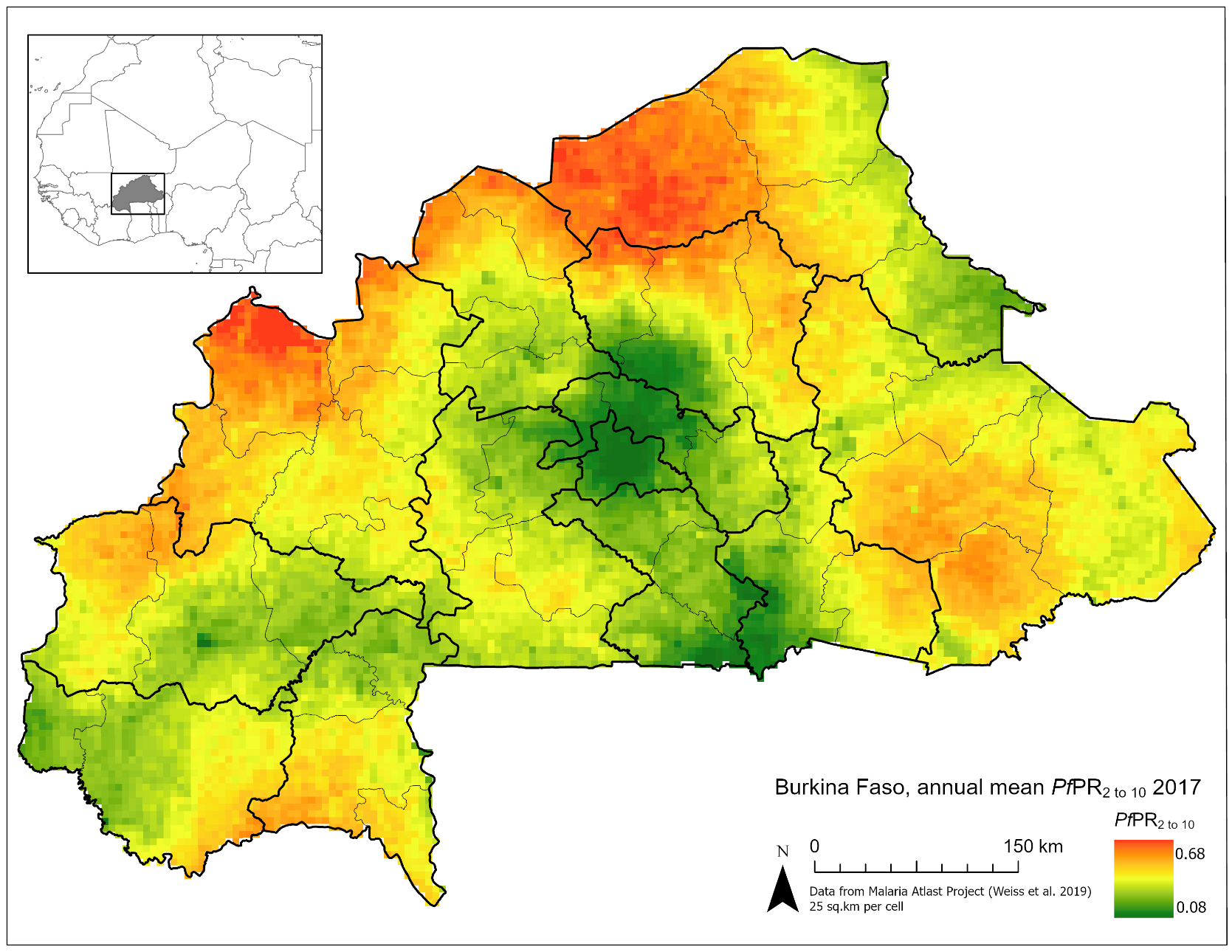

### 4 Figure S3. Climate Zones of Burkina Faso

Climate zones for Burkina Faso determined using data from WorldClim 2.0 (Fick & Hijmans, 2017), boundary data from World Bank Group (2018).

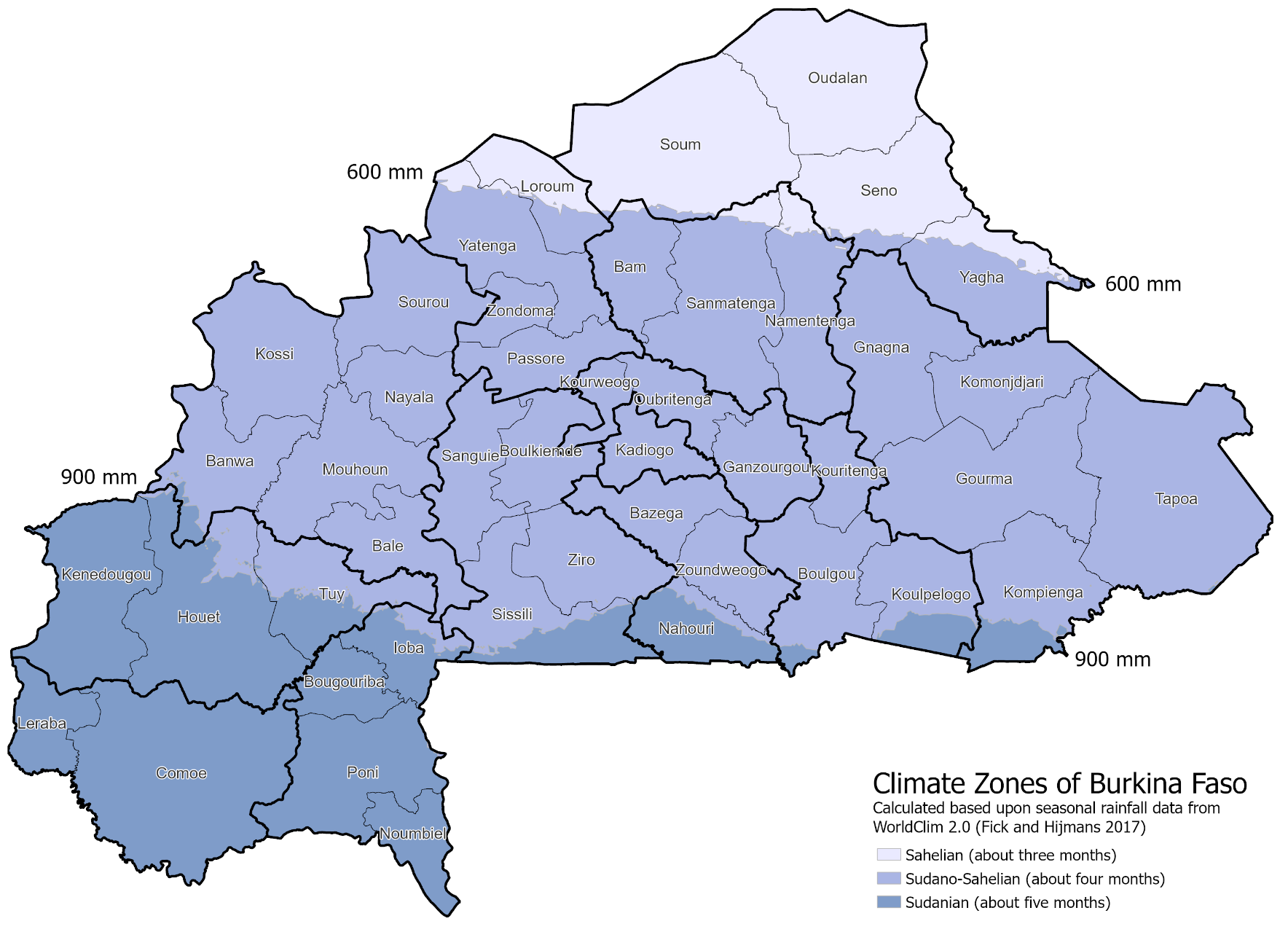

### 5 Figure S4. Probability of importation into a given cell

As calculated using Monte Carlo simulation over one million trials, boundary data from World Bank Group (2018).

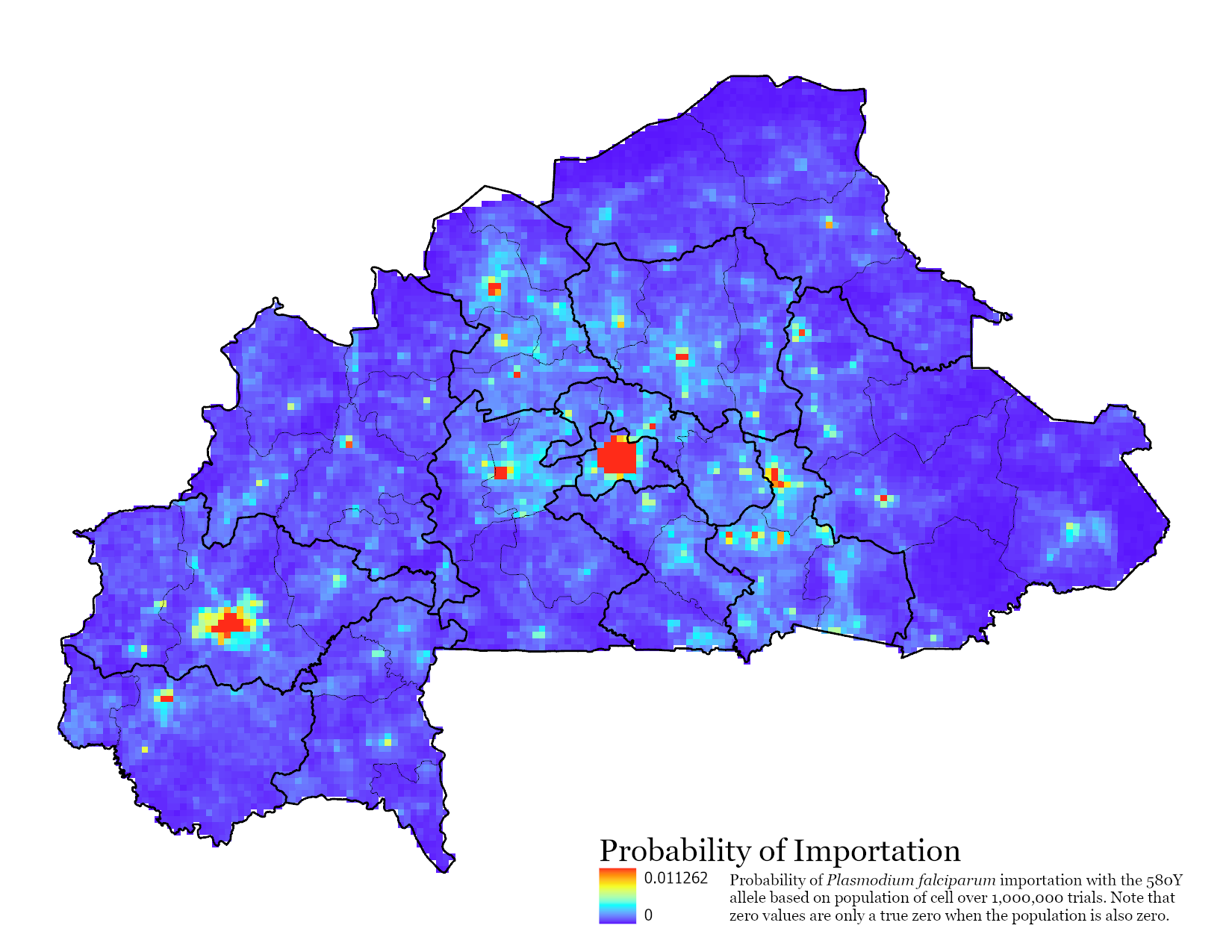
